# Meis2 is required for GABAergic amacrine cell survival in the developing retina

**DOI:** 10.64898/2026.10.06.757052

**Authors:** Shaherah Alqahtani, Patrick C. Kerstein

## Abstract

Amacrine cells (ACs) are the most diverse neuron in the retina, with more than sixty different subtypes, each contributing to the detection of specific visual features. Despite their importance in visual feature detection, the genetic basis for this diversity is still not well understood. Recent single-cell transcriptomic datasets have identified several potential genes that may regulate cellular diversity, but the underlying mechanisms have not been investigated. For example, the transcription factor *Meis2* is selectively expressed in the GABAergic subclass of ACs. In this study, we focused on how *Meis2* regulates the development of GABAergic ACs. We hypothesized that *Meis2* is required for specification and survival of GABAergic ACs. To test this, we used the Meis2 conditional knockout mouse (*Meis2^CKO^*) where *Meis2* was deleted from either the entire retina or specifically from AC precursor cells. The deletion of Meis2 led to a ∼60% reduction in total GABAergic ACs in the retina. This loss of GABAergic ACs occurred only after postnatal day 5 (P5), suggesting *Meis2* is not required for early specification but may be required for survival. To further investigate cell death and survival, we quantified cleaved caspase-3 positive cells in the retina at several developmental timepoints. In *Meis2^CKO^*mice, we found an increase in cleaved caspase-3 immunolabeling at P5 and P6 timepoints coinciding with the reduction of GABAergic ACs. Further analysis of GABAergic AC subtypes demonstrated a subtype-specific cell death in *Meis2^CKO^*retinas. For example, the total number of dopaminergic (TH⁺) ACs were significantly reduced, whereas starburst (ChAT⁺) ACs remained unchanged. These findings suggest that *Meis2* is important for the survival of a subset of GABAergic AC subtypes in the mammalian retina. In addition, *Meis2* may control the expression of specific survival factors that prevent apoptosis during developmental cell death in the retina.

## 1. Introduction

The neural retina is essential for the detection of light and visual features in our environment. During development, proper vision depends on the generation of the correct retinal cell types and the precise assembly into functional neural circuits. All retinal neurons differentiate from a common pool of retinal progenitor cells; a process that requires tightly regulated gene expression to specify each neuronal subtype. However, the genes that control retinal differentiation remain poorly understood. Defining these genetic factors is important for understanding the mechanisms the guide retinal development.

The mature retina contains more than 120 cell subtypes across five major cell classes, each with their own important function during visual processing. Of the major cell classes, inhibitory interneurons called amacrine cells (ACs) exhibit the greatest diversity in cell subtypes morphologically and molecularly. Early morphological studies identified about 45 distinct AC subtypes, whereas single cell transcriptomic analysis has identified at least 63 molecularly distinct AC subtypes (Helmstaedter et al., 2013; Li et al., 2024; Yan et al., 2020). In addition, a recent transcriptomic study from 24 different species have identified at least 43 orthologous AC subtypes that are conserved among vertebrates (Tommasini et al., 2026). Each AC subtype has a unique morphology and function necessary for the detection of a single visual feature, and the loss or dysfunction of a single AC subtype will lead to changes in vision (Calbiague-Garcia et al., 2025; Morrie & Feller, 2016). Importantly, the selective degeneration and dysfunction of AC subtypes is observed in and may contribute to common visual disorders, such as glaucoma and diabetic retinopathy (Eggers, 2023; Gastinger et al., 2006; Gramlich et al., 2021). Despite their importance, the mechanisms that drive AC subtype diversification during development are poorly understood. Identifying these mechanisms will provide insight into molecular determinants of AC identity, their contribution to vision, and potential pathways necessary for retinal repair and regeneration.

Most AC subtypes are broadly categorized into two subclasses based on their neurotransmitter expression, morphology, and function (Calbiague-Garcia et al., 2025). First, the GABAergic ACs have wide, mono-stratified dendrites that contribute functionally to spatial and directional information within retinal circuits. Second, the Glycinergic ACs have narrow, multi-stratified dendrites that are important for vertical communication between retinal layers and play a key role in processes such as dark adaptation. These AC subclasses are born during distinct developmental time windows, with the majority of GABAergic ACs born between embryonic day 12 (E12) and E17, while most Glycinergic ACs are born between E15 and postnatal day 1 (P1)(Cherry et al., 2009; Voinescu et al., 2009). Despite these differences in mature and developing ACs, the genes that control the distinction between GABAergic and Glycinergic ACs are unknown.

Transcription factors play a central role in AC development by regulating their cell fate, differentiation, survival, and connectivity. Some of these transcription factors regulate the specification of the entire AC lineage, while others act more selectively to define individual AC subtypes. Transcription factors, such as *Foxn4* and *Ptf1a*, are required for the specification of the entire AC lineage. The deletion of these transcription factors leads to the loss of ACs and an increase in other retinal neuron cell fates, such as retinal ganglion cells (Fujitani et al., 2006; Li et al., 2004; Nakhai et al., 2007). Other transcription factors act more selectively to control the development of specific AC subtypes. For example, *NeuroD6*, *Isl1*, *Fezf1*, and *Gbx2*, are necessary for the development and survival of single AC subtypes, but not for broad classes of ACs (Elshatory et al., 2007; Kay et al., 2011; Kerstein et al., 2020; Khandokar et al., 2026). However, the transcription factors that govern the distinction to GABAergic and Glycinergic ACs are unknown. Previous studies have demonstrated that the transcription factors, *Bhlhe22* and *Barhl2* are necessary for subsets of GABAergic and Glycinergic AC lineages (Feng et al., 2006; Mo et al., 2004). Recent single cell transcriptomic studies have shed light on this question and have identified the expression of *Meis2* and *Tcf4* as transcription factors selectively enriched in GABAergic and Glycinergic ACs, respectively (Yan et al., 2020). These data suggest *Meis2* and *Tcf4* may contribute to the subclass-specific development of ACs.

Here, we investigated the role of *Meis2* during the development of GABAergic ACs. *Meis2* encodes for a homeodomain transcription factor required for multiple developmental processes, including cardiac morphogenesis, neural crest formation, and nervous system development (Machon et al., 2015). In humans, loss-of-function variants of *MEIS2* are associated with cardiac deficits, cranial-facial anomalies, and intellectual disabilities (Giliberti et al., 2020; Johansson et al., 2014; Louw et al., 2015). In mice, the deletion of *Meis2* is embryonically lethal, however, the recent development of *Meis2* conditional knockout mice enabled the investigation of *Meis2* in neuronal populations and structures that mature later in development (Su et al., 2022; Yang et al., 2021). For example, *Meis2* is required for the differentiation of GABAergic projection neurons in both the striatum and forebrain (Dvoretskova et al., 2024; Yang et al., 2021). The deletion of *Meis2* from these interneurons leads to a selective loss of a subset of medium spiny neurons in the striatum (Su et al., 2022). In the retina, *Meis1* and *Meis2* function together to pattern the early optic cup and promote the generation of multipotent retinal progenitor cells (Dupacova et al., 2021). Furthermore, overexpression of these factors in retinal progenitor cells changes the temporal identity of progenitor cells, biasing them toward early-born neuronal cell fates (Leavey et al., 2025). Although the roles of *Meis1* and *Meis2* in postmitotic retinal neurons remain unclear, several studies have demonstrated the selective expression of *Meis2* in the GABAergic ACs, suggesting a potential role in their development and function (Bumsted-O’Brien et al., 2007; Yan et al., 2020).

In the present study, we used the *Meis2* conditional knockout mouse model to investigate the role of *Meis2* in the developing GABAergic ACs. We observed the abundance of *Meis2* protein is low in the early postnatal retina but begins to increase starting at P4 specifically within the GABAergic ACs. The deletion of *Meis2* from the entire retina or from AC precursors resulted in a significant reduction in the total number of ACs, with a pronounced reduction in the GABAergic population. Furthermore, we observed this decrease occurred after P4 suggesting these ACs were generated normally but fail to survive developmental cell death. Consistent with this, we detected an increase cleaved caspase-3 labeling in ACs at P5, suggesting an increase in apoptosis within the AC population. Together, this study shows that *Meis2* is not required for the early specification of GABAergic AC fate but is necessary for their survival during postnatal retinal development.

## 2. Materials and methods

### 2.1. Animals and Animal Procedures

All experiments were performed using mice of both sexes and all mice were maintained on a C57BL/6J background. The following Cre mouse lines were used for tissue-specific gene manipulation: *Ptf1a*^Cre^ (Nakhai et al., 2007) and *Six3*^Cre^ (Furuta et al., 2000), and *Meis2^Flox/Flox^*(Machon et al., 2015). All mice were housed in a temperature-controlled environment on a 12:12 hour light: dark cycle with unrestricted access to food and water. Offspring of all breeding pairs were genotyped by PCR with primers specific for *Meis2* or *Cre* recombinase. All animal procedures were approved by the Purdue Animal Care and Use Committee (PACUC) at Purdue University and were conducted in accordance with the NIH Guide for the Care and Use of Laboratory Animals.

### 2.2. Immunohistochemistry

Neonatal and adult retinas were prepared for immunolabeling by removing the eyes from the heads of a recently euthanized mice. For tissue fixation, the corneas were removed, and the eye cups were placed in 4% EM-grade paraformaldehyde (PFA) for 30 mins at room temperature. Next, the eyes were washed 3 times in 1X Phosphate-Buffered Saline (PBS) for 10 mins. Eyes were cryo-preserved by placing them in 15% sucrose for at least 2 hours at 4°C. Next, the lenses were removed and the eye cups were embedded in a cryo-mold in Optimal Cutting Temperature media and frozen. Using a cryostat, 20 µm retinal sections were collected on a glass microscope slide (Fisher Superfrost Plus). For immunolabeling, retinal sections were washed in 1X PBS for 10 mins, and then blocked with 2% normal donkey serum (NDS) with 0.2% Triton X-100 for 30 mins. Retinal sections were incubated with primary antibodies in NDS blocking buffer overnight at 4°C. The primary antibodies and their dilutions used in this study are listed in Table 1. Next, sections were washed 3 times in NDS blocking buffer for 10 mins and incubated in secondary antibodies in blocking buffer for 2 hours at room temperature. Retinal sections were washed three times in PBS for 10 mins with DAPI (1:5000) stain included in the first wash step. Tissue was mounted in Fluoromount-G (Southern Biotech) and cover-slipped for imaging.

**Table 1.**

| Antibody | Concentration | Source | Identifier |
| --- | --- | --- | --- |
| Mouse anti-Meis2 | 1:200 | Developmental Studies Hybridoma Bank | Cat#: PCRP-MEIS2-2B3 |
| Rabbit anti-RBPMS | 1:500 | Fisher Scientific (Millipore Sigma) | Cat#: ABN1362MI |
| Mouse anti-Chx10 | 1:200 | Santa Cruz Biotech | Cat#: sc-365519 |
| Rabbit anti-Calbindin | 1:2000 | Swant | Cat#: CB38 |
| Rabbit anti-GlyT1 | 1:1000 | Synaptic Systems | Cat#: 272 103 |
| Goat Anti-ChAT | 1:500 | Fisher Scientific (MilliporeSigma) | Cat#: AB144P200UL |
| Rabbit Anti-TFAP2α | 1:500 | Abcam | Cat#: ab108311 |
| Mouse Anti-GAD67 | 1:500 | Fisher Scientific (MilliporeSigma) | Cat#: MAB5406MI |
| Rabbit Anti-Tyrosine Hydroxylase | 1:500 | Fisher Scientific (Millipore) | Cat#: AB152MI |
| Invitrogen™ Isolectin GS-IB4 Alexa Fluor™ 647 Conjugate | 1:500 | Fisher Scientific | Cat#: I32450 |
| Alexa Fluor® 488 Donkey anti-Mouse IgG (H+L) Antibody | 1:500 | Fisher Scientific (Life Technologies) | Cat#: A21202 |
| Alexa Fluor® 488 Donkey anti-Goat IgG (H+L) Antibody | 1:500 | Fisher Scientific (Life Technologies) | Cat#: A11055 |
| Alexa Fluor® 546 Donkey anti-Rabbit IgG (H+L) Antibody | 1:500 | Fisher Scientific (Life Technologies) | Cat#: A10040 |

For retinal flat-mounts, eyes were dissected and washed as described above. The retinas were removed from the eye cup and flattened by making 4 equally spaced incisions into the center of the tissue. Retinas were washed 3 times in 2% NDS and 0.2% Triton X-100 for 30 mins at room temperature. Retinas were incubated with primary antibodies in blocking solution for 7 days at 4°C or room temperature. After incubation with the primary antibody, the retinas were washed 3 times with blocking solution for 30 mins. Next, retinas were incubated with a secondary antibody in blocking solution overnight at 4°C. Retinas were then washed three times in PBS for 30 mins with DAPI (1:5000) stain included in the first wash step. Tissue was mounted on a slide with Fluoromount-G and cover-slipped for imaging.

### 2.3. Fluorescence image acquisition

All retinas were imaged on a Zeiss LSM 900 laser-scanning confocal microscope using Zeiss Zen software. For retinal sections, immunolabeled cells were capture using a 20X objective. For retinal flat mounts, images were captured either with a 20X objective for cell density measurements or 10X objective for whole retina images. For whole retina images, multiple images were stitched together using the Zeiss Zen software. For all immunolabeling experiments, 6 different images were acquired per animal for off-line analysis.

### 2.4. Fluorescence image analysis

All images were analyzed off-line using the FIJI software(Schindelin et al., 2012). Cell density measurements were made in either retinal flatmounts or cryosections. In retinal flat-mounts, cell density was quantified from a 200×200μm region that was at least 500μm from the optic nerve head and the retina edge. For each animal 4 measurements were made from the central and peripheral retina. In retinal sections, cell density was quantified across a 200μm length of the retina. for each animal, 4 measurements were made across the retinal section at least 500μm from the optic nerve head and retinal edge. Quantification of total ACs using TFAP2α antibody was conducted by measuring the thickness of TFAF2α layer divided by total retina thickness.

### 2.5. Statistical analysis

In these analyses, each experiment included measurements from a minimum of 3 retinas from at least 3 different animals. For all datasets, the measurements and variance were reported as mean ± SEM. Each dataset was first tested for normality with a D’Agostino & Pearson test. For analysis between two groups statistical significance was determined using an unpaired student’s t-test. For analysis between multiple groups (≥3 groups), we used one-way analysis of variance (ANOVA) with a Tukey’s multiple comparisons test. For analysis between multiple timepoints and genotypes a two-way ANOVA with a Tukey’s multiple comparison test was used. All statistical tests were performed using Prism 10 software (GraphPad Software, Inc.)

## 3. Results

### 3.1. Meis2 is expressed in ACs in the postnatal mouse retina

To determine the temporal pattern of *Meis2* expression in the retina, we analyzed retinal sections immunolabeled for *Meis2* from wild-type mice at several postnatal timepoints (P1–P14). We did not observe *Meis2* expression in P1 retinas (Fig. 1A-B), but we began to detect *Meis2* in the INL and GCL by P4 (Fig. 1C-D). The total number of *Meis2*^+^ cells increased between P4 and P5 and remained stable until P14 (n=3 mice p<0.05; Two-way ANOVA with Tukey’s post hoc test) (Fig. 1E–M). Furthermore, *Meis2* co-localized with the TFAP2α+ ACs in the INL and GCL (Fig. 1D, F, H, J, L). These findings indicate that *Meis2* expression in ACs begins at P4 in retinal ACs and remains stable throughout postnatal retinal development.

**Figure 1.**
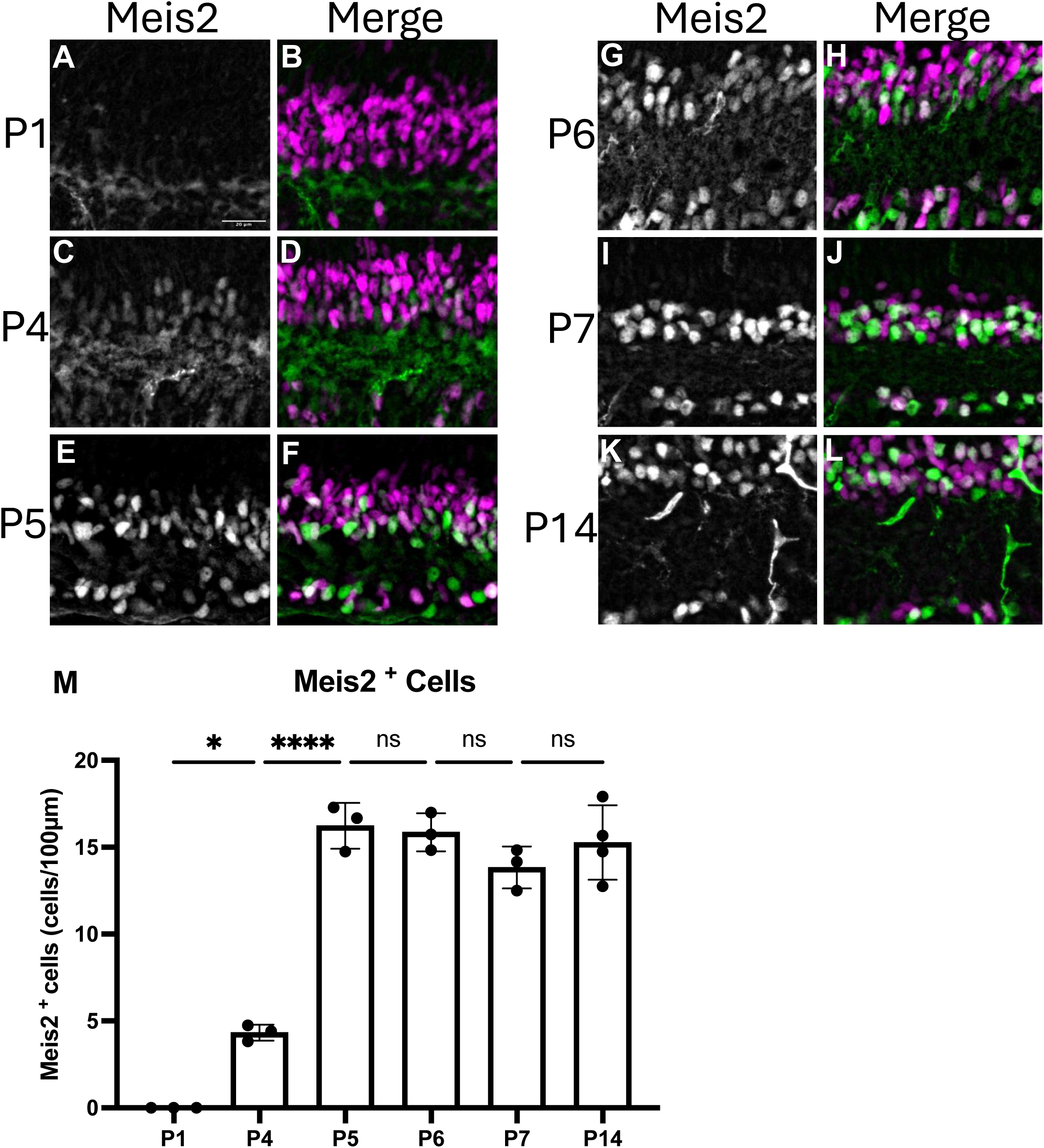
Developmental expression pattern of *Meis2* in the postnatal retina. (A-L) Immunolabeling of the inner retina with *Meis2* (green) and *TFAP2* (magenta) in the wild-type retina during postnatal development (P1–P14). Retinas were immunolabeled with Meis2 (left) and merged with AC marker *TFAP2* (right) at (A-B) P1, (C-D) P4, (E-F) P5, (G-H) P6, (I-J) P7, and (K-L) P14 developmental timepoints. (M) Quantification of the total number of *Meis2+* cells per 100 µm of retinal length within the INL at each developmental stage. Data are presented as mean ± SEM (n = 3 mice per timepoint). Statistical analysis was performed using One-way ANOVA with Tukey’s multiple comparison test. *p < 0.05, ****p < 0.0001. Scale bar: 20 µm (A).

### 3.2. Developmental deletion of *Meis2* leads to a reduction in ACs and IPL thickness

To determine whether *Meis2* is required for AC development, we generated two different *Meis2* conditional knockout mice. First, we used *Six3^Cre^*; *Meis2^Flox/Flox^* mice to delete *Meis2* from the majority of the developing retina (Furuta et al., 2000). Second, we used *Ptf1a^Cre^*; *Meis2^Flox/Flox^* mice to delete *Meis2* from the majority of AC precursor cells (Nakhai et al., 2007). To quantify the total number of ACs we immunolabeled retinal sections for AC marker TFAP2α in P14 retinas from both *Six3^Cre^*; *Meis2^Flox/Flox^*and *Ptf1a^Cre^*; *Meis2^Flox/Flox^* mutant mice. In *Six3^Cre^*; *Meis2^Flox/Flox^* retinas, we observed around a 41% reduction in the total number of TFAP2α+ cells compared to heterozygous control retinas (Fig. 2C–E) We also observed around 24% decrease in IPL thickness in *Six3^Cre^*; *Meis2^Flox/Flox^* retinas (Fig. 2C-D, F) (n=4 mice/genotype, p<0.05; t-test). Similarly, when we deleted *Meis2* from AC precursor cells in Ptf*1a*^Cre^; *Meis2*^Flox/Flox^ retinas, we found both a significant, around 53% reduction of TFAP2α+ cells (Fig. 2G–K) and a thinning of the IPL (Fig. 2L) (n=4 mice/genotype, p<0.05; t-test). Our findings indicate that *Meis2* promotes either the formation or survival of ACs and contributes to IPL organization during retinal development.

**Figure 2.**
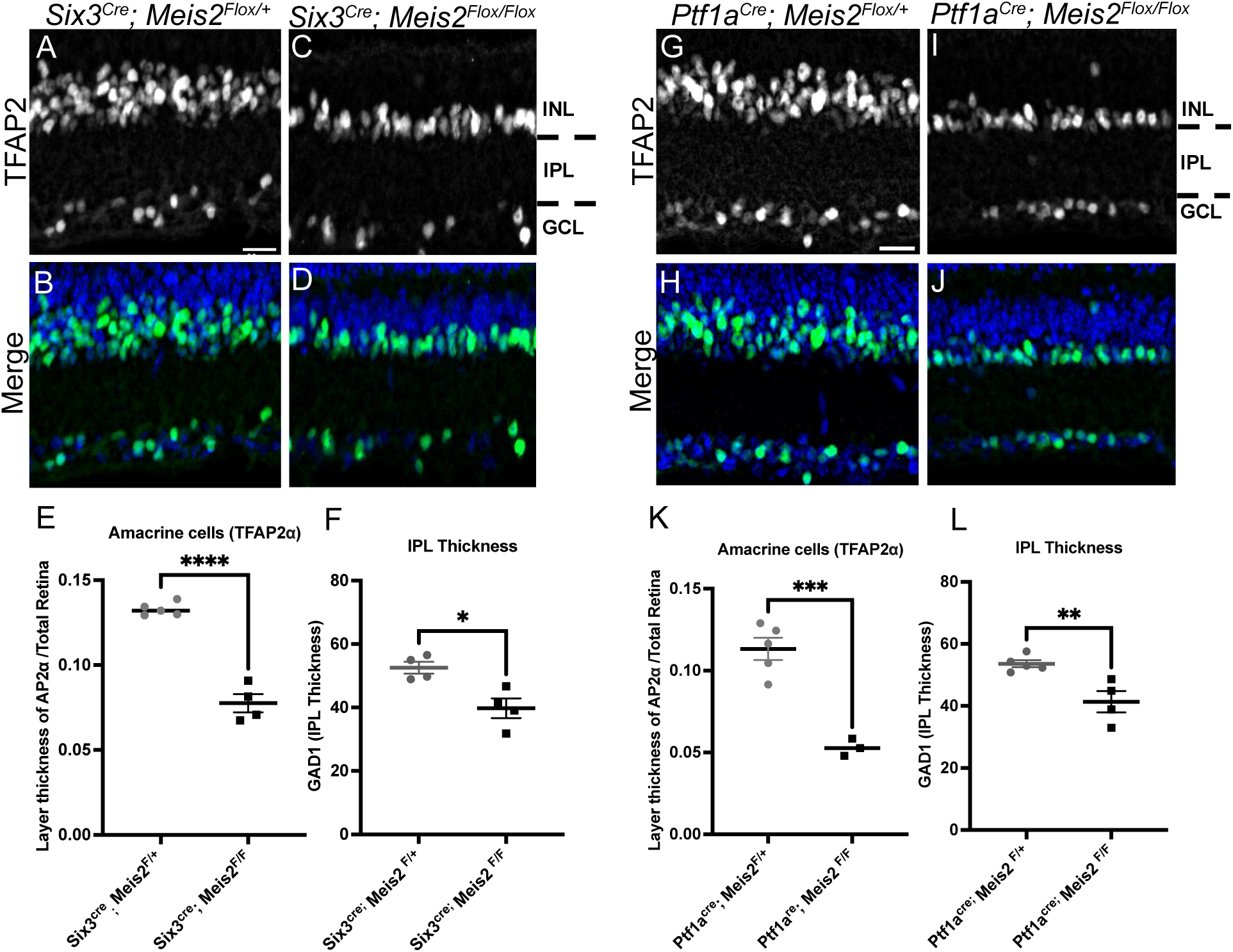
Developmental deletion of Meis2 from the whole retina or amacrine cell precursors reduces amacrine cell density and IPL thickness. (A-D) Cross-sections through the inner retina immunolabeled for (A, B) TFAP2 and (B, D) merged with DAPI (blue) from (A-B) control and (C-D) *Six3*^Cre^; *Meis2*^Flox/Flox^ conditional knockout mice. (E-F) Quantification of (E) TFAP2^+^ AC layer thickness relative to total retinal thickness and (F) IPL thickness in control and Six3^Cre^; *Meis2*^Flox/Flox^ mice. (G-J) Cross-sections from the inner retina of (G, H) control and (I, J) *Ptf1a*^Cre^; *Meis2*^Flox/Flox^ conditional knockout mice immunolabeled for (G-I) TFAP2 and (H-J) merged with DAPI (blue). (K-L) Quantification of (K) TFAP2^+^ AC layer thickness relative to total retinal thickness and (L) IPL thickness in control and Ptf1a^Cre^; *Meis2*^Flox/Flox^ mice. Data represent mean ± SEM from n ≥ 4 mice per genotype. Statistical significance was determined using an unpaired t-test (p < 0.05). *p<0.05, **p<0.01, ***p<0.001, ****p<0.0001. Scale bars: 20 µm (A, G).

### 3.3. Developmental deletion of *Meis2* reduces the density of GABAergic ACs

Previous studies have shown that *Meis2* is highly expressed in GABAergic ACs compared to the Glycinergic AC population (Bumsted-O’Brien et al., 2007). Here, we hypothesized that *Meis2* is required for the selective development of GABAergic ACs. To test our hypothesis, we deleted *Meis2* from the developing retina (*Six3^Cre^*) or AC precursors (*Ptf1a^Cre^*), as described above. In P14 retinal sections, we immunolabeled for GAD1, a marker of GABAergic ACs, and quantified the total GAD1+ cells in the INL of control and *Meis2* mutant retinas. When *Meis2* was deleted across the developing retina, we observed ∼36% reduction in the total number of GAD1^+^ cells in *Six3^Cre^*; *Meis2^Flox/Flox^* retinas compared to heterozygous controls (n=4 mice/genotype, p<0.05; t-test) (Fig. 3A–F, M). Similarly, the selective deletion of *Meis2* in AC precursors resulted in a significant ∼53% reduction in the total number of GAD1^+^ cells in *Ptf1a*^Cre^; *Meis2*^Flox/Flox^ retinas compared to control retina (n=4 mice/genotype, p<0.05; t-test) (Fig. 3G–L,N). In both *Six3^Cre^*; *Meis2^Flox/Flox^* and *Ptf1a*^Cre^; *Meis2*^Flox/Flox^ mutant mice, we observed no changes in the total number of Glycinergic ACs (Supplemental Figure 2A-C). These findings demonstrate that *Meis2* is required for either the generation or survival of a subset of GABAergic ACs during postnatal retinal development.

**Figure 3.**
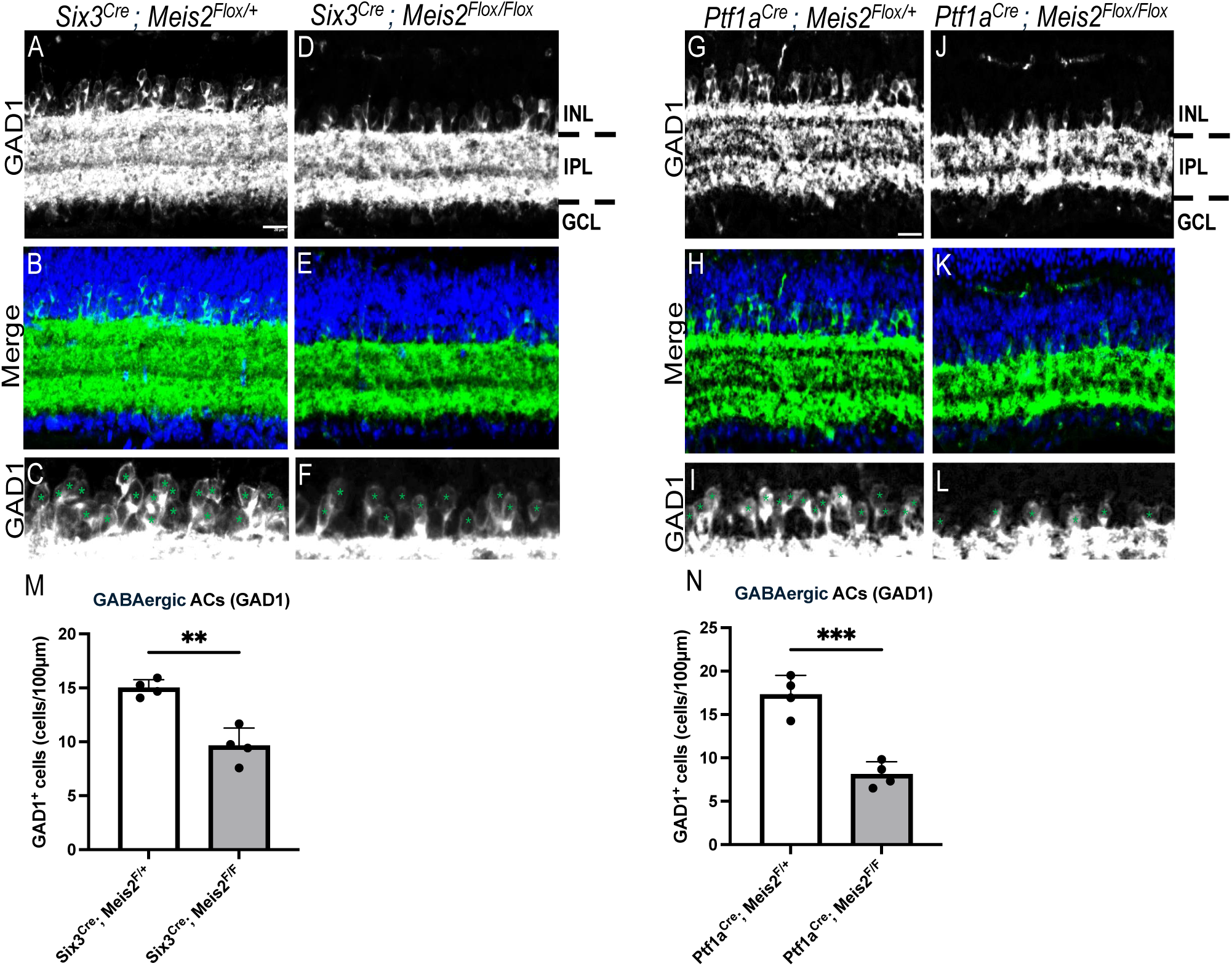
Developmental deletion of *Meis2* reduces the cell density of GABAergic ACs. (A-F) Retinal cross-sections immunolabeled for (A, D) GAD1 and (B, E) merged with DAPI (blue) from (A-C) control and (D-F) *Six3*^Cre^; Meis2^Flox/Flox^ conditional knockout mice. (C, F) Magnified image of the GAD1^+^ cell bodies (asterisks) in the INL from (A) and (D). (M) Quantification of total GAD1^+^ GABAergic ACs per 100 µm of retinal length in control and *Six3*^Cre^; *Meis2*^Flox/Flox^ mice. (G-L) Retinal cross-sections immunolabled for (G, J) GAD1 and (H, K) merged with DAPI (blue) from (G-I) control and (J-L) Ptf1a^Cre^; *Meis2*^Flox/Flox^ conditional knockout mice. (I, L) Magnified image of the GAD1^+^ cell bodies (asterisks) in the INL from (G) and (J). (N) Quantification of GAD1^+^ GABAergic ACs per 100 µm of retinal length in control and *Six3*^Cre^; *Meis2*^Flox/Flox^ mice. Data represented as mean ± SEM from n ≥ 4 mice per genotype. Statistical significance was determined using an unpaired t-test (p < 0.05). **p<0.01. Scale bars: 20 µm in (A, G).

### 3.4. Developmental deletion of *Meis2* reduces the total number of dopaminergic ACs but not cholinergic (ChAT⁺) ACs

Our data show that *Meis2* is required in only a subset of GABAergic ACs. To further examine whether *Meis2* differentially affects specific GABAergic AC subtypes we investigated the effects of *Meis2* deletion on the dopaminergic and starburst AC subtypes. Previous work has reported high *Meis2* expression in TH+ dopaminergic ACs and low expression in ChAT+ Starburst ACs (Yan et al., 2020). In *Six3*^Cre^, *Meis2*^Flox/+^ controls, TH+ cells were present in both the central and peripheral retina (Fig. 4A-B, G). In contrast, *Six3*^Cre^; *Meis2*^Flox/Flox^ retinas exhibited a complete absence of TH+ cells in central, but not peripheral retina (Fig. 4C-D, G). By comparison, the density of ChAT+ starburst ACs—including populations in the INL and GCL—did not differ between heterozygous control and *Six3*^Cre^; *Meis2*^Flox/Flox^ mutant retinas in either the central or peripheral retina (n=3 mice/genotype, p<0.05; Two-way ANOVA with Tukey’s post hoc test) (Fig. 4E-F, H). In contrast, the deletion of *Meis2* from AC precursor cells (*Ptf1a*^Cre^; *Meis2*^Flox/Flox^) exhibited a near-complete loss of TH+ cells in both central and peripheral (Fig. 4I-L, O). However, the density of ChAT+ Starburst ACs were unchanged between the *Ptf1a*^Cre^; *Meis2*^Flox/Flox^ mutant and control retinas (n=3 mice/genotype, p<0.05; Two-way ANOVA with Tukey’s post hoc test) (Fig. 4M-N, P). Together, these results demonstrate that *Meis2* is required for the generation of TH+ dopaminergic ACs, whereas ChAT+ starburst ACs are unaffected, indicating a subtype-specific dependence for *Meis2* in GABAergic ACs.

**Figure 4.**
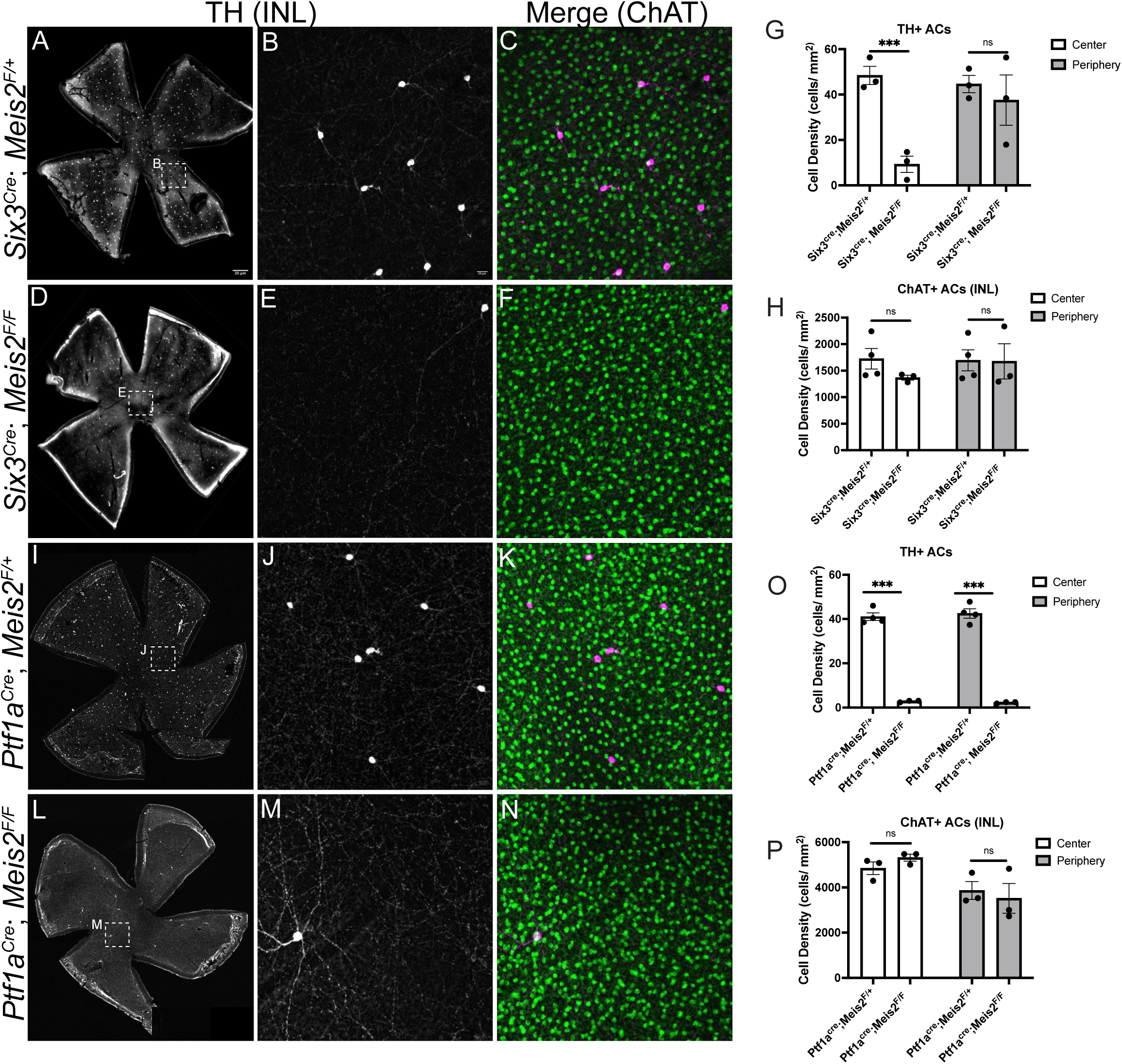
Developmental deletion of *Meis2* reduces the density of dopaminergic (TH⁺) but not cholinergic (ChAT⁺) ACs. (A-D) Retinal flatmounts immunolabeled for TH⁺ cells in (A-B) control and (D-E) Six3^Cre^; *Meis2*^Flox/Flox^ mice. (C-F) Immunolabeling for ChAT⁺ cells in retinal flatmounts from (C) control and (F) *Six3*^Cre^; *Meis2*^Flox/Flox^ mice. (G-H) Quantification of (G) TH⁺ and (H) ChAT⁺ cell density in the central and peripheral retina (n = 3 mice per genotype; p < 0.05, Two-Way ANOVA with Bonferroni Correction). (I-L) Retinal flatmounts immunolabeled for TH⁺ cells in (I-J) control and (L-M) *Ptf1a*^Cre^; *Meis2*^Flox/Flox^ mice. (K-N) Immunolabeling for ChAT⁺ cells in retinal flatmounts from (I) control and (L) Ptf1a^Cre^; *Meis2*^Flox/Flox^ mice. (O-P) Quantification of (O) TH⁺ and (P) ChAT⁺ cell density in the central and peripheral retinal flatmounts (n= 3 mouse per genotype. Scale bars in (A) and (I), 200 µm and (A,B) 50 µm

### 3.5. GABAergic ACs are generated in the correct numbers but fail to survive in Meis2 mutant mice

To determine when Meis2 is essential for AC development, we quantified the number TFAP2α+ and GAD1+ cells across early postnatal stages in Ptf1a^Cre^; Meis2^Flox/Flox^ mutant mice. At P4, we observed no difference in the total number of TFAP2α+ cells between Ptf1a^Cre^; Meis2^Flox/Flox^ mutant and control retinas (P4) (Fig. 5A-B, E). However, by P7, there was a clear reduction in TFAP2α+ cells compared in Meis2 mutant retinas (Fig. 5C-D, E). Similarly, we did not observe any difference in the density of GAD1+ cells in P4 retinas, but by P7, we observed a significant reduction in the density of GAD1+ cells in Meis2 mutant retinas compared to controls (n = 3 mice genotype/timepoint. p < 0.05; two-way ANOVA with Tukey’s post hoc test) (Fig. 5F-I, J). While there was no difference in GABAergic ACs at P4, there was a significant reduction in this AC population by P7 in Meis2 mutant retinas. These data suggest Meis2 is not required for the generation, but for the survival of GABAergic ACs in the developing retina.

**Figure 5.**
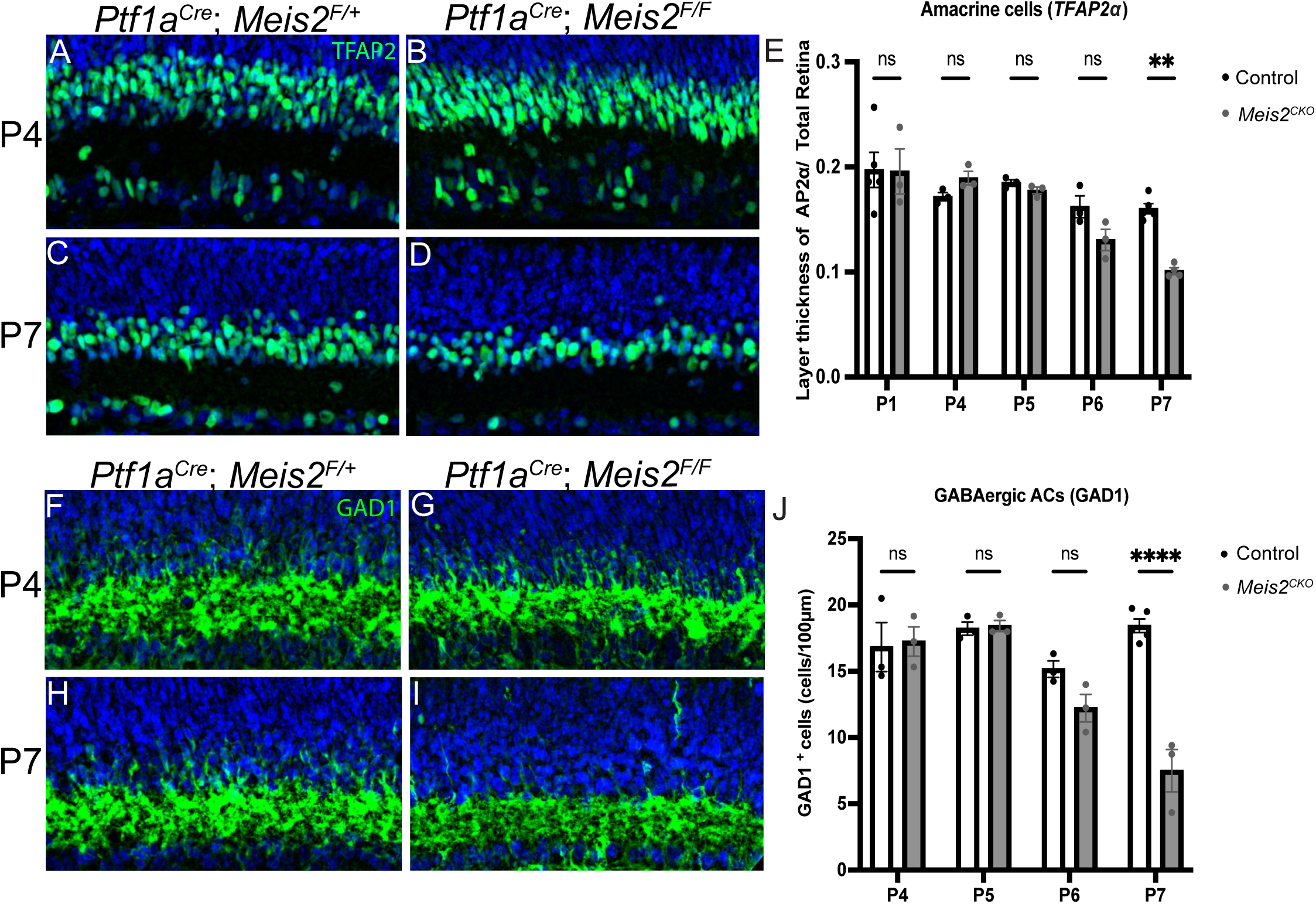
Developmental deletion of *Meis2* reduces the number of GABAergic ACs by postnatal day 7 (P7). (A-D) Retinal sections immunolabeled with AC marker TFAP2a at (A-B) P4 and (C-D) P7 retinas from (A, C) control or (B, D) *Ptf1a*^Cre^; *Meis2*^Flox/Flox^ mice. (E) Quantification of TFAP2a layer thickness relative to total retinal thickness across postnatal time (n ≥ 3 per genotype/timepoint,). (F-I) Retinal sections immunolabeled for GABAergic cell marker GAD1 at (F-G) P4 and (H-I)) P7 from (F, H) control or (G, I) Ptf1a^Cre^; *Meis2*^Flox/Flox^ mice. (J) Quantification of GABAergic AC density from P4 to P7 (n = 3 mice per genotype/timepoint). Quantification (P1–P14) is shown in (E), and the density of GAD1⁺ GABAergic ACs over the same time is shown in (J). Data are presented as mean ± SEM. Statistical significance was determined using two-way ANOVA with Tukey’s post hoc test. **p<0.01; ****p<0.0001; Scale bar: 20 µm (A, F).

### 3.6. Developmental loss of *Meis2* increases apoptosis in the inner nuclear layer

With the deletion of *Meis2*, our data show a loss of GABAergic ACs by P7 in the developing retina. Next, to assess whether the reduction in ACs is linked to increased cell death, we examined apoptosis by immunolabeling for cleaved caspase-3 (CC3) across early postnatal stages. At P4, CC3^+^ cells appeared at similar levels in control and *Ptf1a*^Cre^; *Meis2*^Flox/Flox^ retinas (Fig. 6A-B, E). However, by P5, CC3^+^ cells were significantly increased in *Meis2* mutant mice (n = 3 mice genotype/timepoint. p < 0.05; two-way ANOVA with Tukey’s post hoc test) (Fig. 6C-D, E). We continued to observe a significant increase in CC3+ cells through P6 (n = 3 mice genotype/timepoint. p < 0.05; two-way ANOVA with Tukey’s post hoc test) (Fig. 6E). We further quantified the density of CC3+ cells by retinal layer at P5. At this timepoint, we observed a significant increase in CC3+ cells only in the INL but observed no change in the GCL and ONL of *Meis2* mutant retinas (Fig. 6F). Together, these results show that the loss of *Meis2* leads to increased apoptosis specifically within the INL during postnatal development. Furthermore, this increase in apoptosis precedes the period when AC density begins to decline in *Meis2* mutant retinas.

**Figure 6.**
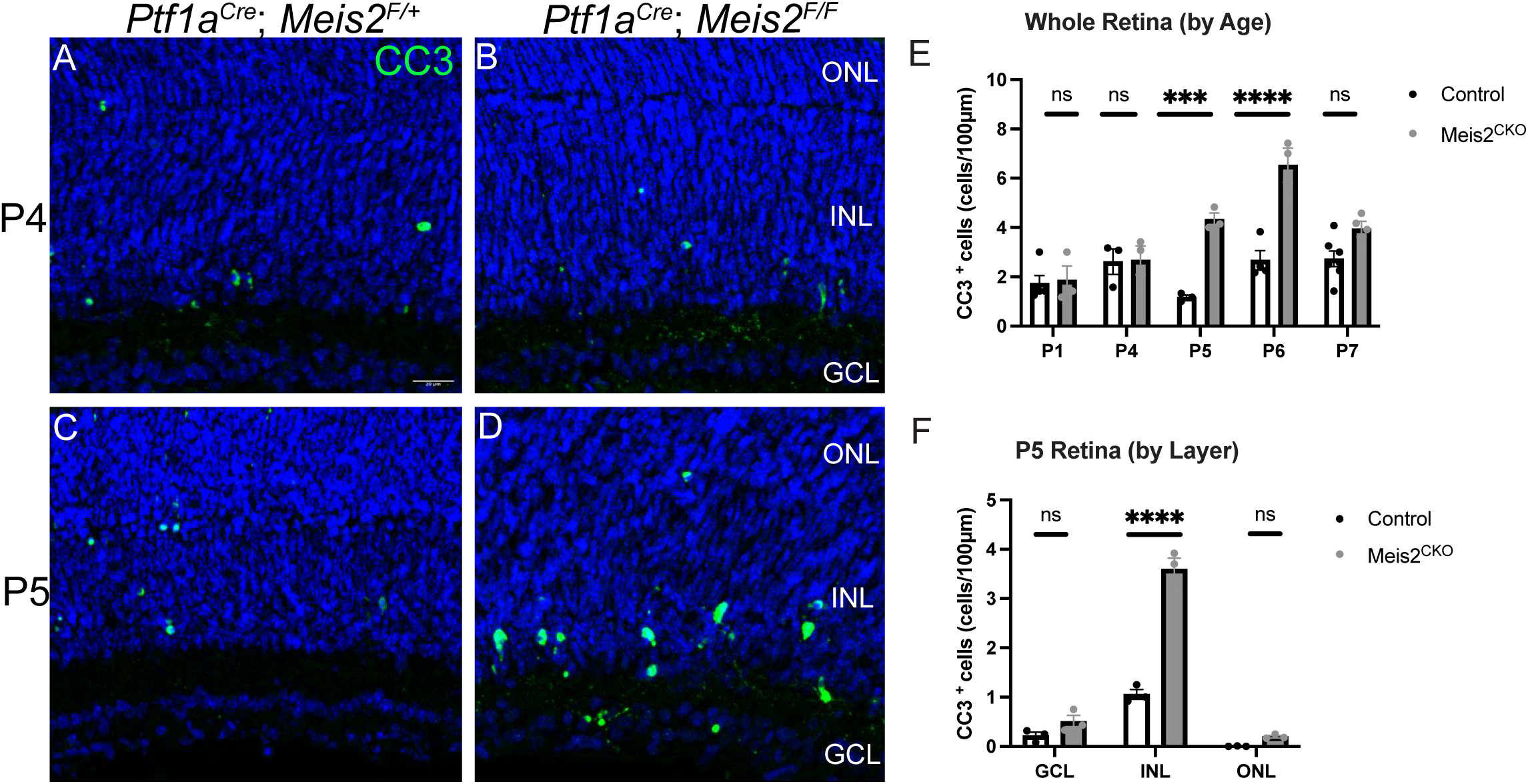
Developmental deletion of *Meis2* increases apoptosis in the inner nuclear layer of P5 retinas. (A-D) Retinal sections immunolabeled for apoptotic cell marker cleaved caspase 3 (CC3) at (A-B) P4 and (C-D) P5 from (A,C) control and (B, D) *Ptf1a*^Cre^; *Meis2*^Flox/Flox^ mice. (E) Quantification of CC3+ cells100 µm of retinal length within the INL at each developmental stage from P1 to P7 ages. (F) Quantification of CC3+ cells at P5 retina separated by retinal location. Data are presented as mean ± SEM (n = 3 mice per genotype/timepoint). Statistical significance was determined using two-way ANOVA with Tukey’s post hoc test. *p<0.05; **p<0.01; ***p<0.001. Scale bar: 20 µm (A).

## 4. Discussion

Neuronal development requires the precise coordination of cell fate specification, differentiation, and survival. Disruption of any of these processes results in selective cell loss and neural circuit dysfunction. In this study, we identified *Meis2* as a critical postnatal survival factor for GABAergic ACs in the developing retina. We found that *Meis2* is expressed at low levels during early postnatal development, however its expression increases in TFAP2α+ ACs starting around P4 (Fig. 1C-D). When we deleted *Meis2* from the entire retina or from AC precursors, we observed a significant reduction in the total number of ACs (Fig. 2C–E and 2G–K). This reduction was due to the loss of only GABAergic ACs (Fig. 3A-N), as the number of Glycinergic ACs were unaffected (Supp. Fig. 2A-C). Importantly, the loss of GABAergic ACs occurred after ACs differentiation suggesting that *Meis2* is not required for cell fate but instead supports cell survival (Fig. 6F-G). Furthermore, shortly after the increase in *Meis2* expression, we observed an increase in cleaved caspase-3 labeling in ACs at P5. Together, these findings demonstrate that *Meis2* plays an essential role in the survival of GABAergic ACs during postnatal retinal development.

The development of inhibitory retinal interneurons, ACs and horizontal cells (HCs), is governed by a transcriptional network that sequentially restricts progenitor competence, promotes differentiation, and specifies inhibitory interneuron fate. Early transcription factors, such as *Foxn4,* act in mitotically active retinal progenitors to bias the lineage output towards ACs and HCs while suppressing alternative photoreceptor and ganglion cell programs (Li et al., 2004). Upon cell-cycle exit, *Ptf1a* promotes a fate-restricted precursor population necessary to specify AC and HC identities, while preventing differentiation into retinal ganglion cells (Fujitani et al., 2006). Downstream of *Ptf1a*, the transcription factors *Tfap2a* and *Tfap2b* act to drive differentiation of all AC subtypes. The loss of *Tfap2a/b* leads to the failure of AC differentiation and a switch to a photoreceptor cell fate (Jin et al., 2015). Downstream of *Tfap2a/b*, it is unclear what transcription factors control differentiation between GABAergic and Glycinergic ACs. Based on recent single cell transcriptomic datasets, expression of the transcription factor *Meis2* is highly restricted to GABAergic ACs (Yan et al., 2020). Based on its expression pattern, we initially hypothesized that *Meis2* would control GABAergic AC specification in the developing retina. However, our results in the present study suggest differently. First, we found *Meis2* abundance is low in the early postnatal retina during AC differentiation but exhibits highest abundance starting at P4 after the differentiation of GABAergic ACs (Fig. 1C-D). Second, the deletion of *Meis2* from the developing AC precursors does not affect the initial cell specification of ACs, as GABAergic ACs appear in normal numbers at P4 (Figure 6 F, G). Only after a wave of developmental cell death at P7 do we observe a reduction in GABAergic ACs (Fig. 6 I). Together, the timing of *Meis2* upregulation with the increase in apoptosis observed suggest Meis2 is required for GABAergic AC survival and not differentiation. Thus, the *Foxn4*–*Ptf1a*–*Tfap2a/b* cascade appears to generate and differentiate inhibitory interneurons, whereas *Meis2* is required to maintain the survival of the GABAergic ACs population during postnatal retinal development.

During retinal development, ACs undergo a significant wave of programmed cell death, which is essential for establishing mature population sizes and refining spatial organization (Young, 1984). In the inner nuclear layer of the mouse retina, this apoptotic peak occurs primarily between postnatal days 3 and 8 (Vecino & Acera, 2015; Vecino et al., 2004). This natural cell loss serves as a fundamental mechanism for the refinement of retinal mosaics of individual AC subtypes. For example, the regularity of the cholinergic starburst AC arrays is significantly enhanced through natural cell loss (Galli-Resta & Novelli, 2000). Similarly, the VGluT3+ AC population utilizes Bax-mediated cell death to transform an initially random distribution into a regular mosaic (Keeley et al., 2021). Interestingly, VGluT3⁺ AC survival is governed by extrinsic factors rather than solely cell-intrinsic programs. For the VGluT3+ AC population, survival and mosaic patterning depend on trophic support from bipolar cell afferents, as selective Bax deletion in bipolar cells results an increase in the number of VGluT3+ ACs and a disruption of their regular spatial distribution (Keeley et al., 2021). Furthermore, the proapoptotic regulator Bax also controls dopaminergic ACs survival, as the deletion of Bax leads to a greater than fourfold increase in dopaminergic AC density (Whitney et al., 2009). In addition to extrinsic survival cues, intrinsic transcription factors are required to support the survival during this refinement period. The transcription factor *Gbx2* is selectively expressed in S3-stratifying ACs and is necessary for both their development and survival. Embryonic deletion of *Gbx2* from *Ptf1a*⁺ precursors leads to a significant reduction, and loss, of S3-*Gbx2*⁺ ACs, along with disruption of their dendritic lamination within sublamina (Khandokar et al., 2026). Likewise, *Prdm13* is not required for early AC genesis but is essential for the survival of Ebf3⁺/calretinin⁺ AC subtypes (Goodson et al., 2018). Our findings suggest a similar mechanism for *Meis2*. Rather than acting at the initial stage of AC production, *Meis2* functions during the postmitotic developmental window, where it is required for the selective survival of GABAergic ACs. Together, these studies support the idea that AC diversity is established not only by early fate specification but also by subtype-specific survival mechanisms that occur during a window developmental cell death.

In this study, our data suggest *Meis2* functions within GABAergic ACs to regulate their postmitotic survival in the developing retina. Several other genes may also serve as survival factors for GABAergic ACs. For example, genetic ablation of nuclear factors *ROR*β*1* and *Nr4a2/Nurr1* both lead to the reduction of the total number of GABAergic ACs (Jiang & Xiang, 2009; Liu et al., 2013). However, both *ROR*β*1* and *Nr4a2* play a clear role in AC specification early in development and may not have a specific role in AC survival. Furthermore, several adhesion molecules are important for the survival of specific subsets of ACs. For example, perturbation of specific clustered protocadherins isoforms has a dose-dependent response on both dendritic organization and survival of ACs (Ing-Esteves et al., 2018). Within this pathway, the PcdhγC4 isoform exhibits a particularly strong pro-survival activity through its cytoplasmic domain (McLeod et al., 2025). This suggests that downstream intracellular signaling from adhesion molecules can direct ACs survival during retinal development. Other adhesion molecules, such as DSCAMs, also couple dendritic organization with developmental cell death (Li et al., 2015). Modulating the activity of DSCAMs, particularly through a gain-of-function mutation, leads to increased cell death of specific AC subtypes. Finally, outside the retina, transcriptional profiling in striatal neurons identified the adenosine receptor *Adora2a* as a potential downstream target of *Meis2* required for neuronal survival. (Yang et al., 2021). In the retina, *Adora2a* is expressed in several retinal cell types, including the ACs (Liu et al., 2026). Like in the striatum, *Meis2* may regulate the expression of *Adora2a* receptors to control the survival of ACs during retinal development. Together, we hypothesize that *Meis2* maintains GABAergic AC survival through the coordination of effector genes, such as adhesion proteins, protocadherins and DSCAMs, and other receptors, such as Adora2a. Future studies, will need to explore these potential effector genes to more completely understand the mechanisms by which *Meis2* controls AC survival.

Here, we demonstrate the role of *Meis2* in retinal development, however it also plays important roles throughout the nervous system and in human disease. For example, investigations into mouse forebrain development from *Meis2* knockout animals has shown important roles for *Meis2* in GABAergic interneuron development. Similar to our findings in the retina, *Meis2* is required for the developmental survival of GABAergic neurons of the striatum (Yang et al., 2021). Other studies have found a role of *Meis2* in generation and specification of specific neuron populations in the mouse, such as the D1 and D2 medium spiny neurons (Su et al., 2022). Finally, *Meis2* also plays an important role in differentiation of GABAergic projection neurons that develop from the ganglionic eminence (Dvoretskova et al., 2024). In human disease, several studies have demonstrated that *Meis2* haploinsufficiency is associated with both intellectual disability and motor delay, indicating that *Meis2* is critical for normal neural development (Johansson et al., 2014; Louw et al., 2015). Finally, although not directly implicating *Meis2*, recent evidence shows that retinal GABAergic signaling is altered in autism further underscoring the clinical importance of maintaining inhibitory retinal circuitry (Huang et al., 2024). Together, these studies across the nervous system suggest that *Meis2* not only regulates neuronal development but also helps maintain inhibitory neuron lineages throughout maturation. Here, our findings extend the function of *Meis2* to the retina, where we demonstrate its critical and specific role in postmitotic survival of GABAergic ACs. Future studies will need to address the downstream mechanisms of *Meis2* and inhibitory interneuron survival, as these mechanisms will have broad implications for both the retina and other areas of the nervous system.

## Supplement figures

**Figure 1: Developmental deletion of *Meis2* from the whole retina did not affect Ganglion, Photoreceptor, Bipolar cells density.** (A-B) Cross-sections through retina immunolabeled for RBPMS (Green)and merged with DAPI (blue) in control (A) and *Six3*^Cre^; *Meis2*^Flox/Flox^ conditional knockout mice (B). (C) Quantification of RBPMS+ cells density. (D-E) Cross-sections through retina immunolabeled for Chx10 (Green) merged with DAPI (blue). In control (D) and *Six3*^Cre^; *Meis2*^Flox/Flox^ conditional knockout mice (E). (F) Quantification of Chx10+ cells density. (G-H) Cross-sections through retina immunolabeled for DAPI (blue) to label outer nuclear layer (ONL) in control (G) and *Six3*^Cre^; *Meis2*^Flox/Flox^ conditional knockout mice (H). (I) Quantification of ONL thickness. Data represent mean ± SEM from n ≥ 3-4 mice per genotype. Statistical significance was determined using an unpaired t-test (p < 0.05). *p<0.05, **p<0.01, ***p<0.001, ****p<0.0001(F, I) and Statistical significance was determined using two-way ANOVA with Tukey’s post hoc test. *p<0.05; **p<0.01; ***p<0.00(C). Scale bars: 20 µm (A).

**Figure 2: Developmental deletion of *Meis2* from the whole retina did not affect Horizontal and Glycinergic ACs cells density.** (A-B) Retinal cross-sections immunolabeled for GlyT1 (green) merged with DAPI (blue) in control (A) and (B) *Six3*^Cre^; *Meis2*^Flox/Flox^ conditional knockout mice. (C) Quantification GlyT1^+^ Glycinergic ACs per 100 µm of retinal length in control and *Six3*^Cre^; *Meis2*^Flox/Flox^ mice. (D-E) Immunolabeling for Calbindin⁺ cells in retinal flat-mounts from (D) control and (E) *Ptf1a*^Cre^; *Meis2*^Flox/Flox^ mice. (F) Quantification of Calbindin⁺ cell density in retina. Data represent mean ± SEM from n ≥ 3-5 mice per genotype. Statistical significance was determined using an unpaired t-test (p < 0.05). *p<0.05, **p<0.01, ***p<0.001, ****p<0.0001(C-F). Scale bars: 20 µm (A-D).

**Figure 3: Developmental deletion of *Meis2* from the whole retina did not affect cholinergic (ChAT⁺) ACs density in Ganglion Cell Layer (GCL).**

(A-B) Immunolabeling for ChAT⁺ cells in retinal flat-mounts from (A) control and (B) *Six3*^Cre^; *Meis2*^Flox/Flox^ mice. (C) Quantification of ChAT⁺ cell density in the central and peripheral retina (n = 3 mice per genotype; p < 0.05, Two-Way ANOVA with Bonferroni Correction. Scale bars: 20 µm (A).

