## Supplementary figures and images for "Meis2 is required for GABAergic amacrine cell survival in the developing retina"

### Supplemental Figure 1

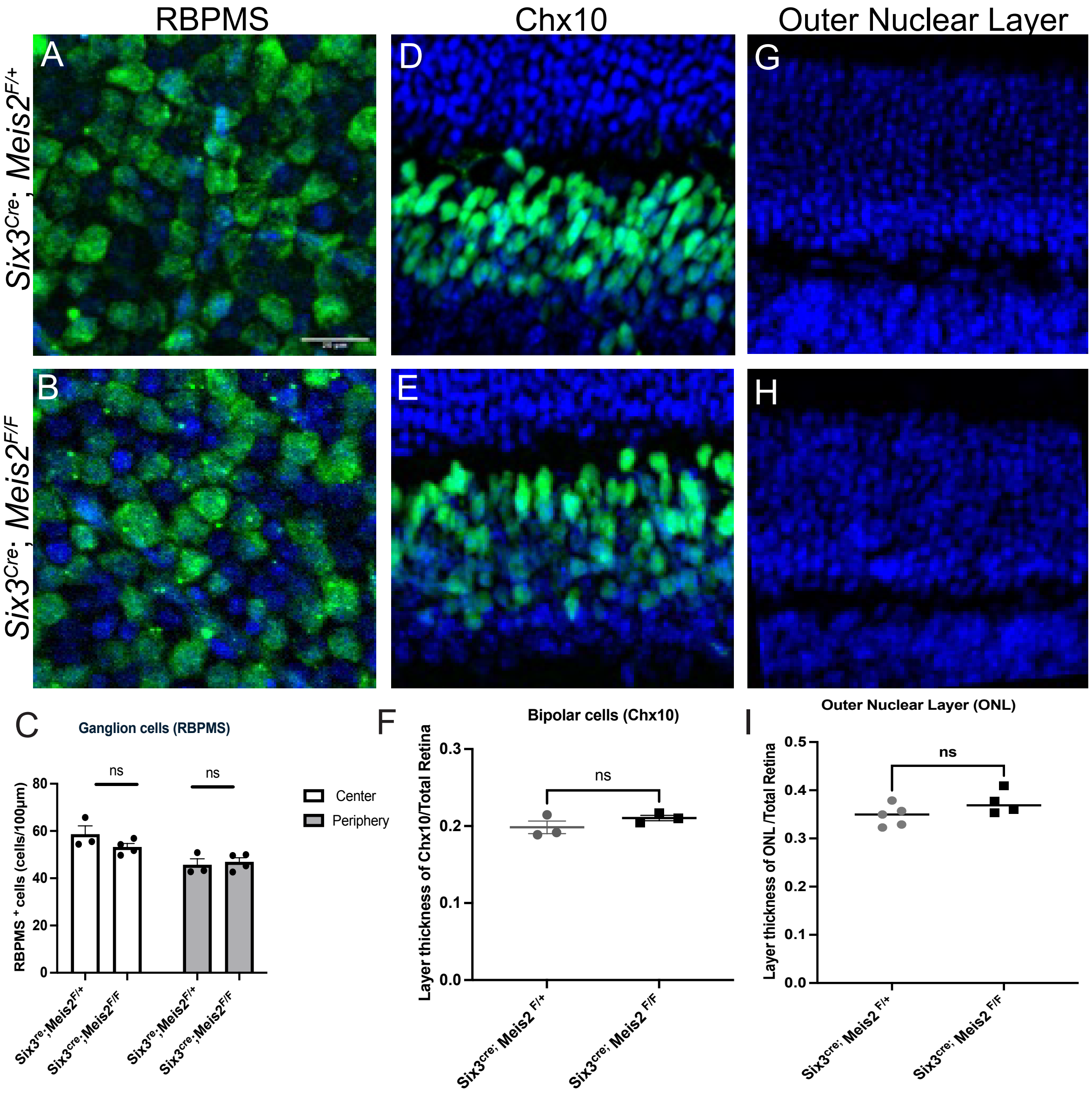

### Supplemental Figure 2

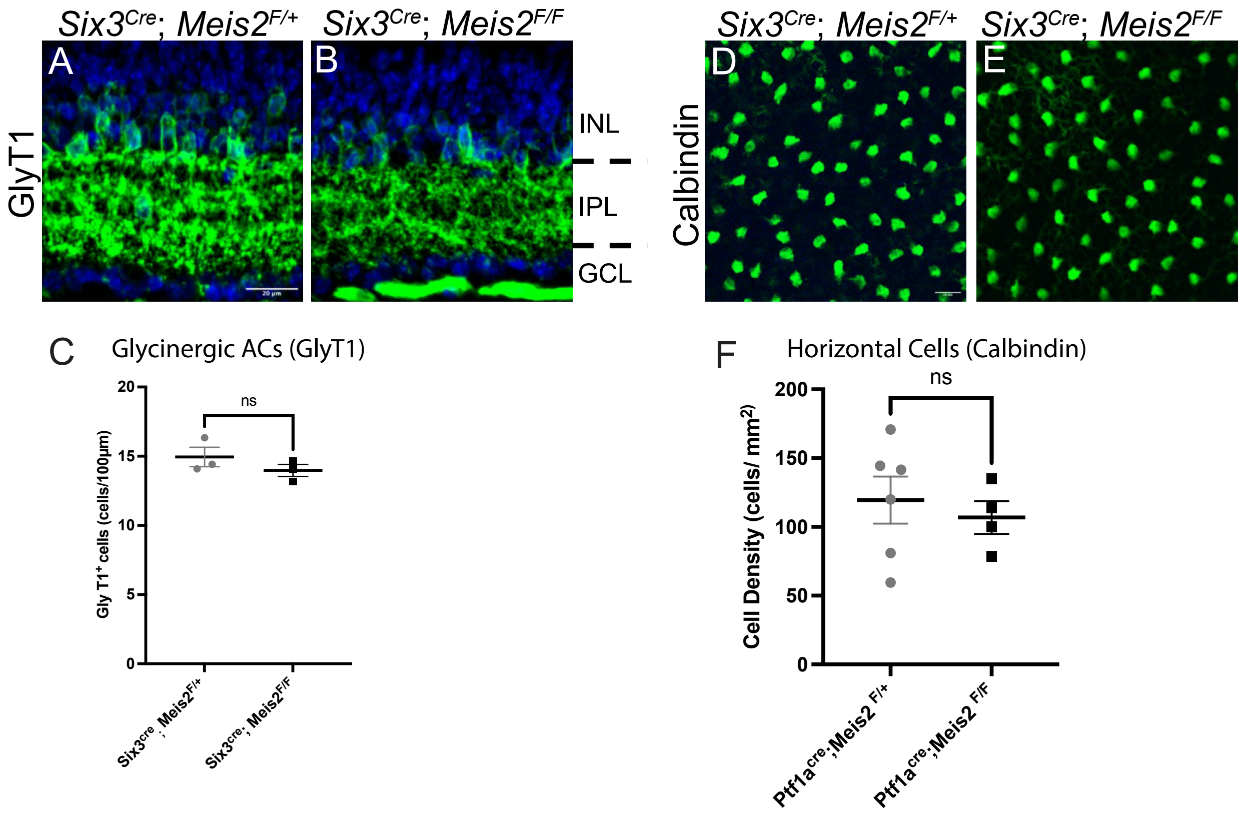

### Supplemental Figure 3

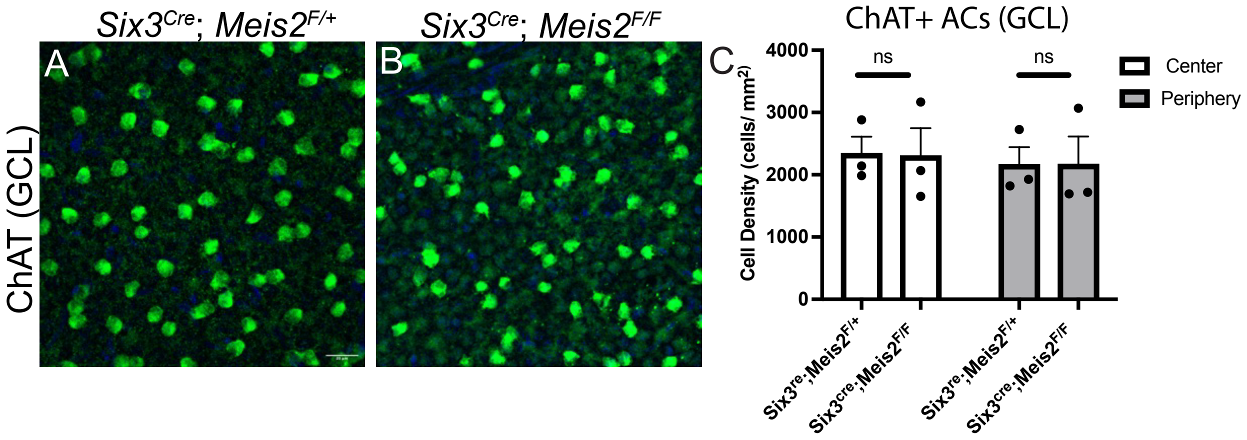
